# A new method to study genome mutations using the information entropy

**DOI:** 10.1101/2021.05.27.445958

**Authors:** Melvin M. Vopson, Samuel C. Robson

## Abstract

We report a non-clinical, mathematical method of studying genetic sequences based on the information theory. Our method involves calculating the information entropy spectrum of genomes by splitting them into “windows” containing a fixed number of nucleotides. The information entropy value of each window is computed using the m-block information entropy formula. We show that the information entropy spectrum of genomes contains sufficient information to allow detection of genetic mutations, as well as possibly predicting future ones. Our study indicates that the best m-block size is 2 and the optimal window size should contain more than 9, and less than 33 nucleotides. In order to implement the proposed technique, we created specialized software, which is freely available. Here we report the successful test of this method on the reference RNA sequence of the SARS-CoV-2 virus collected in Wuhan, Dec. 2019 (MN908947) and one of its randomly selected variants from Taiwan, Feb. 2020 (MT370518), displaying 7 mutations.

## 1. Introduction

Information is a very abstract concept that comes in many forms including analogue information, biologically encoded DNA / RNA information, quantum information and digital information. Shannon developed the classical information theory in 1948 giving the mathematical formulation of the amount of information extracted from observing the occurrence of an event. He is considered the father of modern computing and the inventor of the unit of information, the “bit” [1].

Shannon’s information theory has already underpinned fundamental progress in a diverse range of subjects such as computing [2], cryptography [3], telecommunications [4], physiology [5], linguistics [6], biology [7], geology [8], biochemical signalling [9], mathematics and physics [10-12].

In this article, we report a new methodology based on Shannon’s information theory that allows studying the mutation dynamics in genome sequences and offers a path to predicting future mutations. In fact, using the information theory to study genome sequences is not new. The first reports of analysis of DNA sequences through information theory methods appeared in 1970s. Reichert et al. developed an information-based methodology for determining the quality of an alignment of two code sequences [13]. Other relevant studies include the use of statistical methods for characterizing nucleotidic sequences based on maximum entropy techniques [14], the analysis of repetitive sequences and their effect on the entropy [15] and the study of long-range correlation and complexity in DNA sequences [16-19]. In spite of the successful application of the information theory to the study of genetic sequences, there have been some critical studies, most notable published by Hariri et al., who concluded that the applicability of Shannon’s information theory to genetic sequences did not provide any useful insights into molecular biological sequences [20]. For more recent studies we encourage the readers to consult the review articles published by Vinga [21], Machado [22], as well as a few other new articles on this topic [23-25].

Here, we propose an approach that involves the creation of information entropy (IE) spectra of genome sequences, in order to analyze their mutation dynamics. This approach is applicable to any genome sequence, of any size and it enables new avenues for research in the field of bioinformatics and genetics. The software required to implement this methodology, called GENIES (GENetic Entropy Information Spectrum), has been developed as part of this project and it is freely available to the scientific community [26].

## 2. Information theory for genome studies

Let us assume a set of n independent and distinctive events *X* = {*x*_1_, *x*_2_, …, *x*_*n*_ } having a probability distribution *P* = {*p*_1_, *p*_2_, …, *p*_*n*_ } on X, so that each event x_j_ has a probability of occurring *p* _*j*_ = *p*(*x* _*j*_), where p_j_ ≥ 0 and 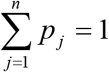 According to Shannon, the average information per event, or the number of bits of information per event, one can extract when observing the set X once is:

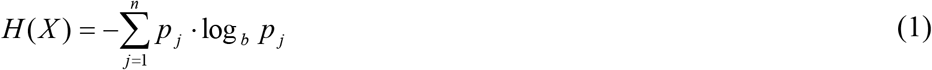

The function *H(X)* resembles an information entropy function and it is maximum when the events xj have equal probabilities of occurring, *p*_*j*_ = 1/ *n*, so *H* _max_ (*x*) = log_*b*_ *n*. When observing N sets of events X, or equivalently observing N times the set of events X, the number of bits of information extracted from the observation is N·H(X). This approach could be used to study other forms of information systems, such as the biologically encoded DNA / RNA information.

Thinking of the genome as a coding system, and given the highly complex nature of genome architecture, the information entropy profile of functional regions of DNA / RNA (e.g. exons, promoters, enhancers, etc.) may represent distinct patterns.

Similarly, the pattern of information entropy would show characteristic changes in response to specific mutations, suggesting a role for information entropy in the determination of genomic variation. Genetic sequences are examined here from an external point of view, as information storage systems, without considering the detailed physical or chemical mechanisms for information processing.

A typical DNA genome sequence can be represented as a linear sequence of the four nucleotides adenine (A), cytosine (C), guanine (G), and thymine (T). For a complete genome, chromosomes represent contiguous sequences, although may contain unknown sequences typically represented by the character N. For simplicity, distinct chromosome sequences can be thought as being part of one continuous string of A, C, G and T letters. Many viruses (including SARS-CoV-2) have RNA genomes, where thymine (T) is replaced by uracil (U), but for simplicity T will be considered for both.

Our set of n independent and distinctive events therefore becomes *X* = {*A,C,G,T*}, with a probability distribution *P* = {*p*_*A*_, *p*_*C*_, *p*_*G*_, *p*_*T*_ }. Using digital information units (b = 2), for four possible distinctive events / states (n = 4) we would require 2 bits per nucleotide (*H* _max_ (*x*) = log_*b*_ *n* = log_2_ 4 = 2) to encode the message: A = 00, C = 01, G = 10, T = 11. Let us consider a random DNA subset, consisting of N = 34 letters:

### CACTTATCATTCTGACTGCTACGGGCAATATGTG

The above subset is randomly generated and the grouping rule of the bases A and T (or U in RNA) and C and G is not respected. If the letters within this DNA subset would have equal probabilities to occur (1/4), then the subset would have H(X) H_max_(X) = 2, and a total entropy of N·H(X) = 68 bits of information. However, the above subset has the following probability distribution:

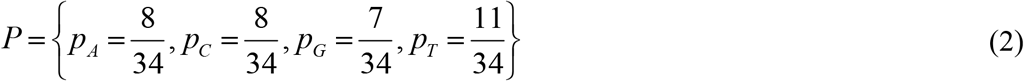

resulting in:

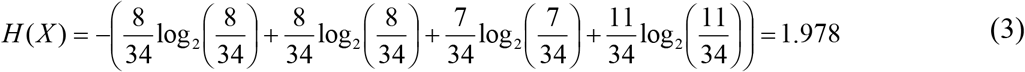

Therefore, instead of 68 bits to encode the sequence we can get by with only 67.25 bits, or the entropy of the subset is 67.25.

Equation (1) and the above approach give information only about the distribution of the individual symbols within the DNA subset and their entropy values, but it doesn’t reflect the correlations between the symbols. In order to incorporate these possible correlations, equation (1) must be generalized to the so-called block information entropies [27]:

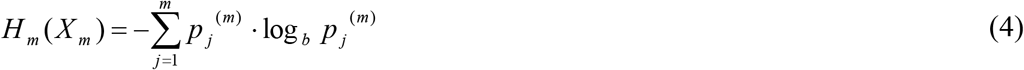

where p^(m)^ are the probabilities of the combinations of m symbols, where 1 ≤ m ≤ n. The index now extends over all possible combinations of m symbols, which are called m-blocks. The block information entropies have been used in numerous studies of the information content of genomic sequences, without targeting mutations [28,29]. In our case m can be any positive integer value larger or equal to 1 and less than or equal to 4. The choice of the number of symbols or m-blocks can be based on the observation that coding sequences for peptides and proteins are encoded via codons, i.e. sequences of blocks of three nucleotides. Hence, choosing m = 3, we will consider each three-nucleotide codon to be a distinct symbol in Shannon’s information theory framework. We can then take a subset of the genome and estimate the probability of occurrence of each codon by simply counting and dividing by the length. Let us consider again the case of our DNA subset, consisting of N = 34 letters:

### CACTTATCATTCTGACTGCTACGGGCAATATGTG

Assuming the readout could start at any nucleotide, counting from left to right each three adjacent nucleotides, a set of 32 three letters combinations with 29 distinct combinations is obtained:

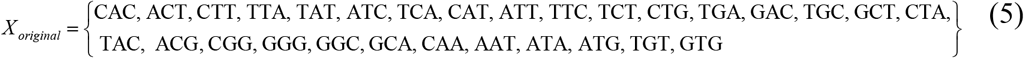

and their probability distribution:

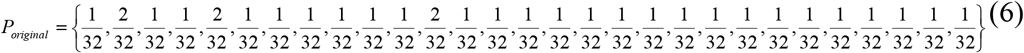

The maximum possible entropy of the DNA subset is log_2_ 29 = 4.858 bits. Using the equation (4) and the probability distribution (6), the actual entropy of the original subset is H_original_ = 4.813 bits.

To demonstrate the principle proposed here for studying genome mutations, let us now assume that the same 34 letters genome subset suffers a random mutation at base point 11, as represented in the figure below:

**Figure 1.**
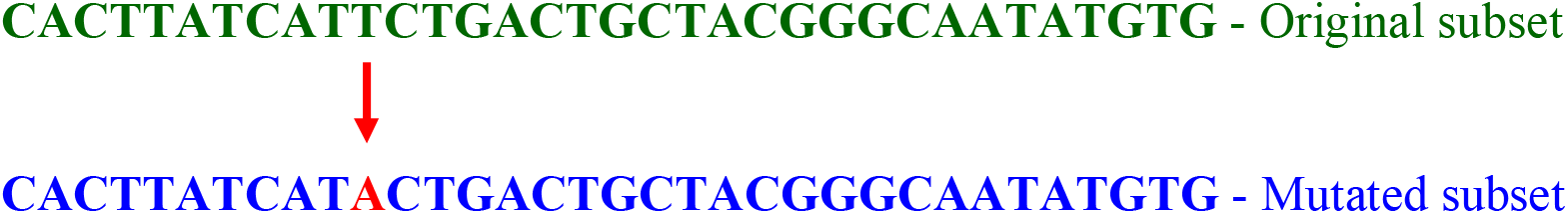
Example of 1 base genome mutation at index 11 (T mutates into A).

The mutated genome subset has again 32 three letters combinations, but 26 distinct instead of 29, as the original subset:

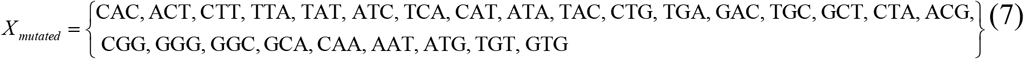

and their probability distribution:

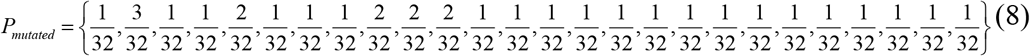

The maximum entropy of the mutated genome subset is log2 26 = 4.7 bits and using equation (4) and the probability distribution (8), the actual entropy of the mutated subset is H_mutated_ = 4.601 bits. This is different to the entropy of the original subset, demonstrating the concept of using the information theory to detect and study mutations. To quantify the difference between the original and the mutated subsets one could use either the difference between the two information entropies, ΔH = H_original_ - H_mutated_ = 0.212 bits, or the Information Entropies Ratio, IER = H_original_ / H_mutated_ = 1.046. Using the difference, any non-zero value would indicate the presence of a mutation within the subset. However, using the ratio, any value not equal to 1 would indicate the occurrence of at least one mutation. From a convenience point of view, the Information Entropy Ratio (IER) is a better representation since the IER parameter has no units, while ΔH has units of bits.

## 3. Information entropy spectrum

Using the concept of information entropy to study genome mutations has been briefly demonstrated in the previous section, for a small genome subset of 34 characters. The main objective is to implement this technique for studying full size genomes. The method proposed here involves computing a so-called Information Entropy (IE) Spectrum of the genome. This is achieved by splitting a large genome into subsets, called “windows”. A “window” has a given number of base points (characters) called “window size”, WS, which is the equivalent of the random genome subset discussed in the previous section, where WS = 34. Starting from left to right, one slides the “window” across the whole genome, where each position of the window is obtained by sliding it for a fixed number of characters, called “step size”, SS. In order to ensure that all sections of the genome are captured by this process, the SS must be at least 1 and maximum WS, so 1< SS ≤ WS. By doing this, a given genome of N characters (number of base points), will result in a total of N_w_ windows, given by the formula:

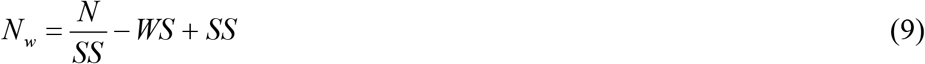

The ratio N/SS is rounded down to the nearest integer value. Since this may result in the last window being incomplete, this incomplete window is ignored. The link between the index of a given i^th^ window and the nucleotide index in the genome sequence corresponding to the first character of the i^th^ window is given by the formula:

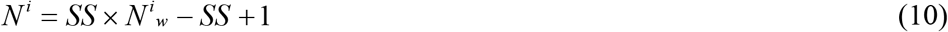

where, *N*^*i*^ *= 1,2,3…N* and *N*^*i*^_*w*_ *= 1,2,3,…N*_*w*_, with N_w_ ≤ N given by relation (9). To study the whole genome, one needs to compute the IE spectrum of the genome, which is obtained by calculating the IE value of each window and plotting the IE values obtained as a function of the window index location within the genome. The window index location in the IE spectrum takes values from 1 to N_w_, depending on the values of N, WS and SS, according to (9). In fact the procedure of window sliding across the whole genome is exactly the same process used to generate the set of 32 three letters combinations of the random 34 bases genome subset in the above example (see relation (5)), where the SS =1 and WS = 3, so N_w_ = 34/1 – 3 + 1 = 32.

To clarify this procedure, we present a graphical example in figure 2, by reconsidering a random genome sequence example. In this example, the window size is WS = 12 and the step size is SS = 4. For these WS and SS parameters, assuming a fictitious genome size of N = 210000 bases, according to (9) we will generate a set of 52492 windows, i.e. N_w_ = 52492. The IE of each window is then computed by counting the distinct m-blocks within each window and their occurrence probabilities. As already mentioned, this was done in the previous example by setting m = 3, so WS = 3 and SS = 1, while N becomes the size of the window itself. According to (9), for each window in this example we get N_w_ = 12/1 – 3 + 1 = 10 in each window containing 3-letter elements. For Window 1, the distinct 3-letter occurrences are: CAC, ACT, CTT, TTA, TAT, ATC, TCA, CAT, ATT, TTC and using equation (4) we get the IE of Window 1, IE_W1_ = 3.322 bits, which is also the maximum possible value as all occurrences have the same probability, 1/10.

**Figure 2.**
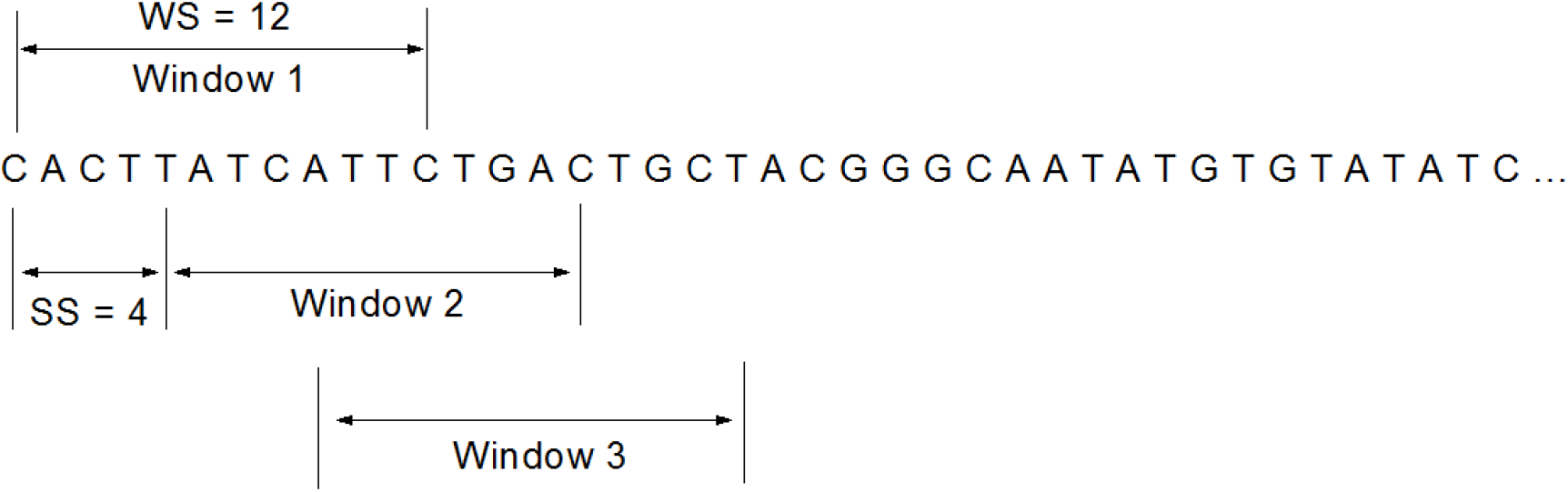
Example of computing the Information Entropy Spectrum of a genome by sliding a window across it, with window size is WS = 12 and step size is SS = 4.

Repeating this process for all 52492 windows, a numerical value for each window is obtained converting the biological information contained within a window to a single numerical value. Plotting these values against their index location within the genome, results in what we call the Information Entropy (IE) Spectrum. The IE spectrum is a numerical representation of the genetically encoded information within a given genome. This is a very convenient algorithm as it allows further processing of the information contained in the spectrum. The correct choice of the m-block size, window size (WS) and step size (SS) is a matter of investigation and it depends on the information that one seeks to extract form the study. In any case, the proposed methodology can only work using fully automated computer software, and we developed LabView software called Genome Information Entropy Spectrum (GENIES) to do this. The program is available free of charge by contacting the corresponding author, or via direct download from the repository [26]. The GENIES user manual can also be downloaded freely [30]. All the operational details of the GENIES software are available in these references [26,30].

## 4. Experimental results

To demonstrate this methodology, we analyzed the reference RNA sequence of the SARS-CoV-2 collected in December 2019 in Wuhan (MN908947) [31], which is currently used as the index reference genome in genomic analyses of the current COVID-19 pandemic. The reference MN908947 sequence contains 29903 nucleotides and is available to download freely from NCBI GenBank database [32]. Figure 3 shows a collection of IE spectra of the reference SARS-CoV-2 sequence MN908947, calculated using GENIES for fixed step size, SS = 3, and variable window size, WS. The IE value of each window was determined using m-block size = 3, with step size = 1 within each window. The data clearly show the WS effect on the IE spectra characteristics. Larger WS values promote larger average IE value of each spectrum, as shown in figures 3 and 4. The maximum IE value per spectrum corresponding to each window size and extracted from the data follows closely the maximum theoretical value expected, log_2_ *n* = log_2_(*WS* − 2), where n is the maximum number of distinct events in a given window, which in our case is n = WS/1 – 3 + 1 = WS – 2. Interestingly, the experimental and the theoretical maximum IE values diverge from each other when WS > 33 (see figure 4). The inflection point at WS = 33 is an important parameter as it gives us a selection criterion of the optimal window size. This is because the ability to extract useful information from the IE spectra requires large changes of information entropy. As data in figure 4 suggests, larger WS values work well, but the maximum IE values are obtained for WS ≤ 33.

**Figure 3.**
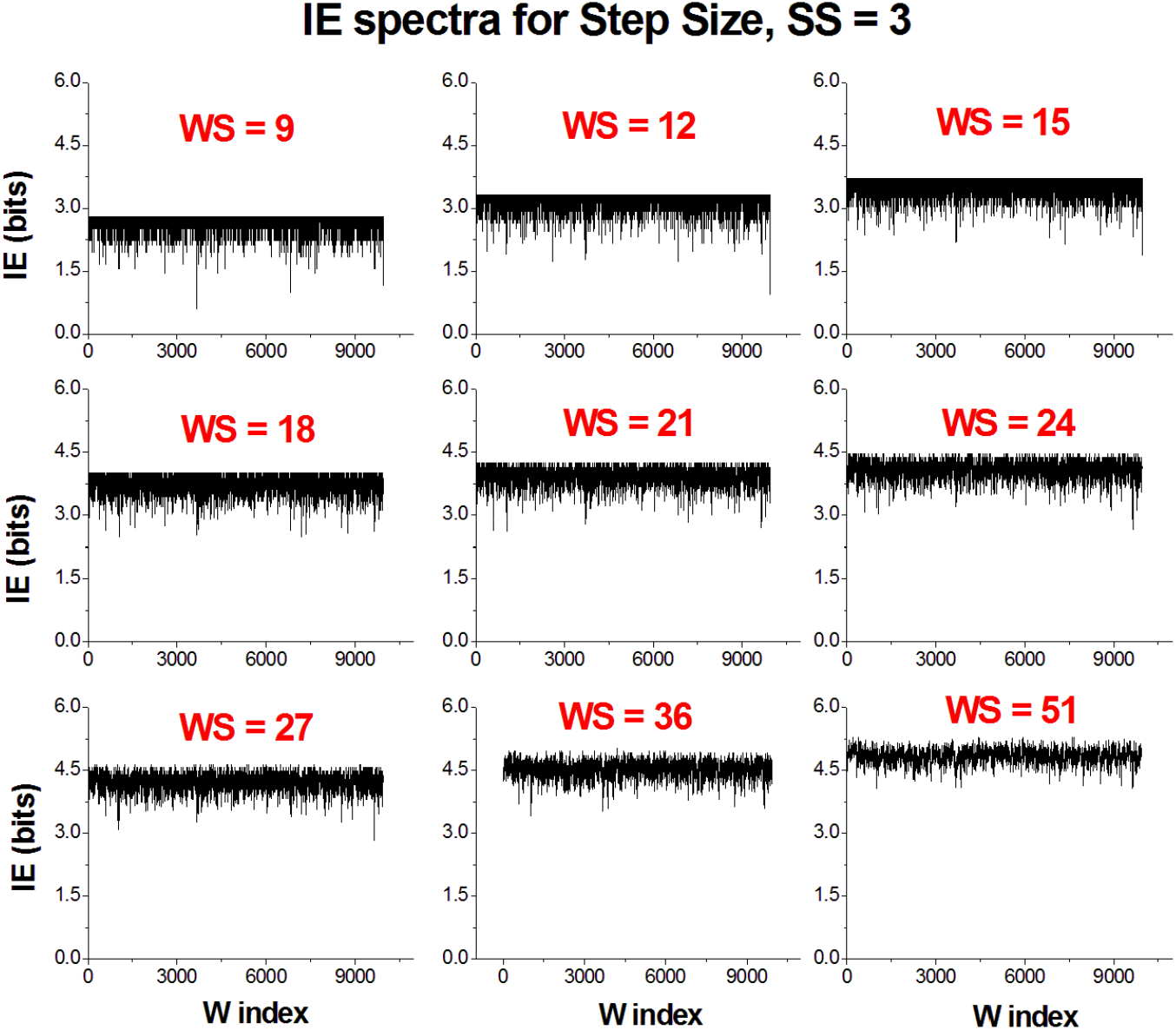
IE spectra of SARS-CoV-2 reference sequence (MN908947) produced using the GENIES software for different window sizes, WS, with fixed step size, SS = 3.

**Figure 4.**
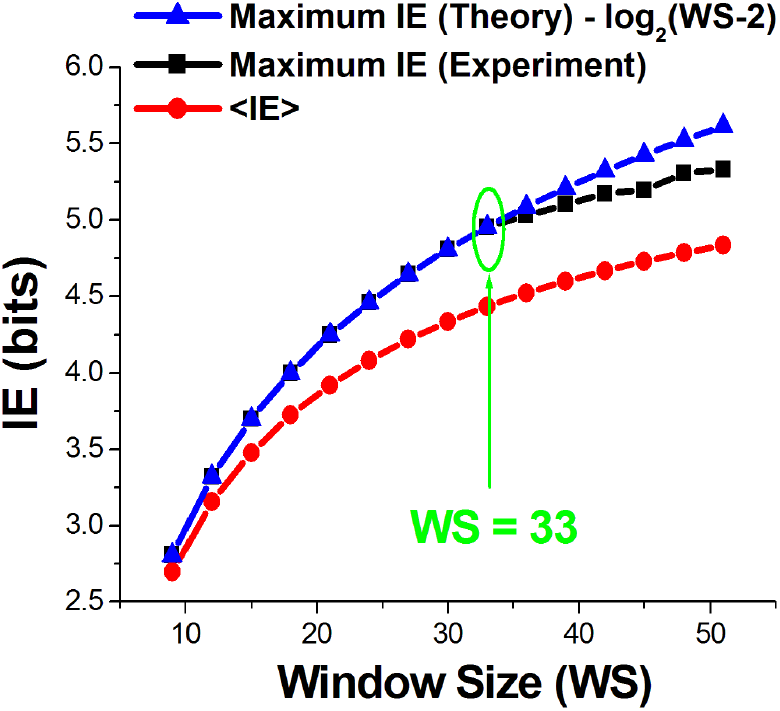
Average IE values, maximum theoretical and experimental IE values per spectrum as a function of WS.

It is important to mention that Thanos, Li and Provata have already reported a similar method to that described here [33]. The key difference between this work and that published in reference [33] is that they divided the sequence into non-overlapping windows called blocks, and they calculated the blocks information entropies by counting the individual nucleotides rather than 3-codon blocks as in our formalism. For a genome size N, the number of windows was N_w_ = N / WS, essentially meaning that the step size was equal to the window size in our formalism, SS = WS in relation (9). For example, taking a random 10000 characters sequence, setting the SS = WS = 100, the approach presented in [33] produces an IE spectrum of N_w_ = 100 size, while our approach with SS = 2, WS = 100 and N = 10000 generates a far more detailed spectrum, capturing possible hidden correlations and having a size 49 times larger, i.e. N_w_ = 4902. Despite this reduction of information, the approach taken by Thanos et al. [33] allowed the extraction of meaningful information related to the detection of repetitive sequences and it was proven very useful in finding evolutionary differences between organisms. Hence, it is expected that the enhanced method proposed here would facilitate additional tools for studying genome sequences, including their mutation dynamics, as demonstrated in the next section.

## 5. Detecting genetic mutations from IER spectra

In section 2 we explained theoretically how the information entropy could facilitate the detection of genetic mutations. In this section we apply this concept to a real genome, by studying the IE spectra of the SARS-CoV-2 reference sequence (MN908947) and one of its randomly selected variants collected in Taiwan on the 27^th^ of February 2020 (MT370518) [34]. If mutations occurred, the IER spectrum obtained should contain values not equal to 1, each indicating a mutation. This technique is a rapid and time effective method of detecting genetic mutations without a full single base point nucleotide comparison between the two genomes. However, for the purpose of testing this method, we first performed a direct comparison of the two genomes and 7 mutations were identified in the MT370518 sequence, as following: C1059T, G1397A, G11083T, C23934T, T28648C, T28688C, G29742T. The first character is the base point that underwent the mutation in the reference genome, the number is its location in the sequence and the second character is what it mutated into in the new sequence.

We computed the IE spectra using the GENIES program under the same conditions described in section 3: m-block size of 3, m-block slide step of 1, window step size SS = 3, and variable window size, WS. Figure 5 shows a collection of IER spectra of the reference sequence (MN908947) divided by one of its variants (MT370518). As expected, for WS ≤ 33 the variations in the IE spectra are more pronounced, resulting in distinctive features in the IER spectra. Unfortunately, the IER spectra failed to reproduce the full set of 7 genetic mutations detected using the direct comparison of the two sequences. Although spectra acquired for different WS values failed to capture all 7 mutations, with best results being 5 - 6 correctly identified mutations, the spectra acquired using smaller WS values captured even fewer mutations. Most notably when WS = 9, only 3 out of 7 mutations were identified. These results correspond to window step size SS = 3. Although not shown here, the same results have been obtained when testing the same procedure for smallest step size, SS = 1, and the largest allowed step size, SS = WS.

**Figure 5.**
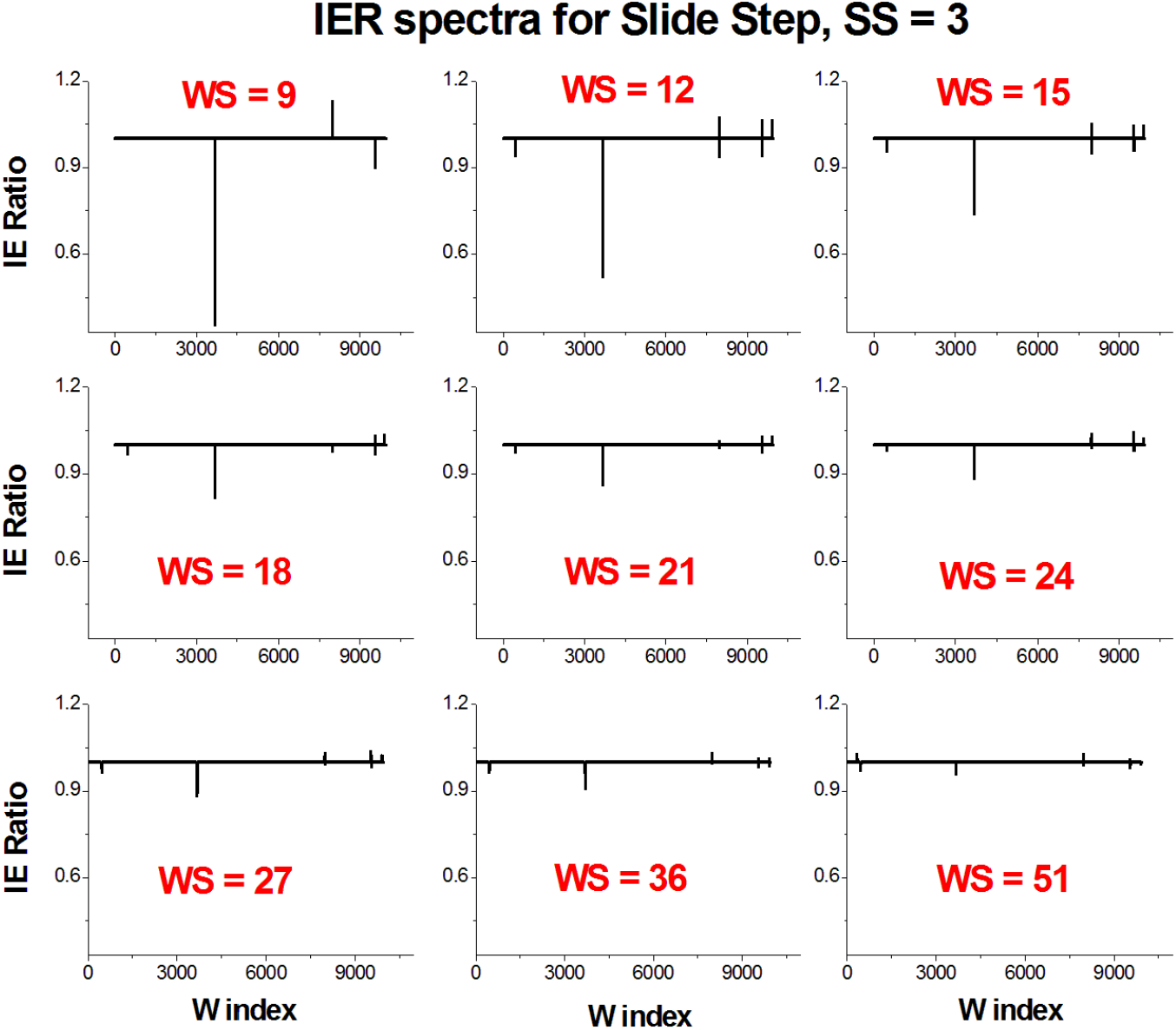
Information entropy ratio (IER) spectra of SARS-CoV-2 reference sequence (MN908947) to its variant (MT370518), produced using the GENIES software for different window sizes, WS, with fixed step size, SS = 3.

Since the IER spectra computed using a range of WS and SS values failed to fully reproduce the genetic mutations, we next turned our attention to investigating the effect of the m-block size. The m-block size can take values from m = 1 to n, where n is the number of distinct events, which in our case is n = 4 corresponding to the set {A, C, G, T}. All our computations so far have been performed for m = 3, but m =1, 2, 4 values are allowed. To investigate the effect of the m size, we fixed the window size to WS = 15 (i.e. following the rule WS ≤ 33) and we chose the step size equal to the window size, SS = WS = 15. A more detailed IE scan of the genome is obtained using SS < WS, but because the windows overlapping, a single mutation could show up as multiple mutations in the IER spectra and further data processing or decomposition is required. When the windows do not overlap, i.e. SS = WS, there is a loss of possible information about correlations between the nucleotides, but the IER spectrum returns the correct number of mutations. Figure 6 shows the results for all possible m-block values, indicating that the larger m-block size reduces the ability to capture the mutations. When m = 4 we obtained 2 mutations out of 7, m = 3 captured 3 mutations, and m = 2 and 1 captured 7 out of 7 mutations. The m = 1 case corresponds to the standard information entropy case as given by relation (1) and it is not a true m-block. However, m = 2 is a more convenient choice as it captures not only all the mutations, but also possible nucleotide-to-nucleotide correlations. The IER spectra in figure 6 are plotted using identical scales to allow direct visual comparison, indicating that m = 2 produces larger, more distinctive features in the IER spectrum than m = 1. The m = 2 appears to be universally valid regardless of the choice of other parameters such as WS and SS.

**Figure 6.**
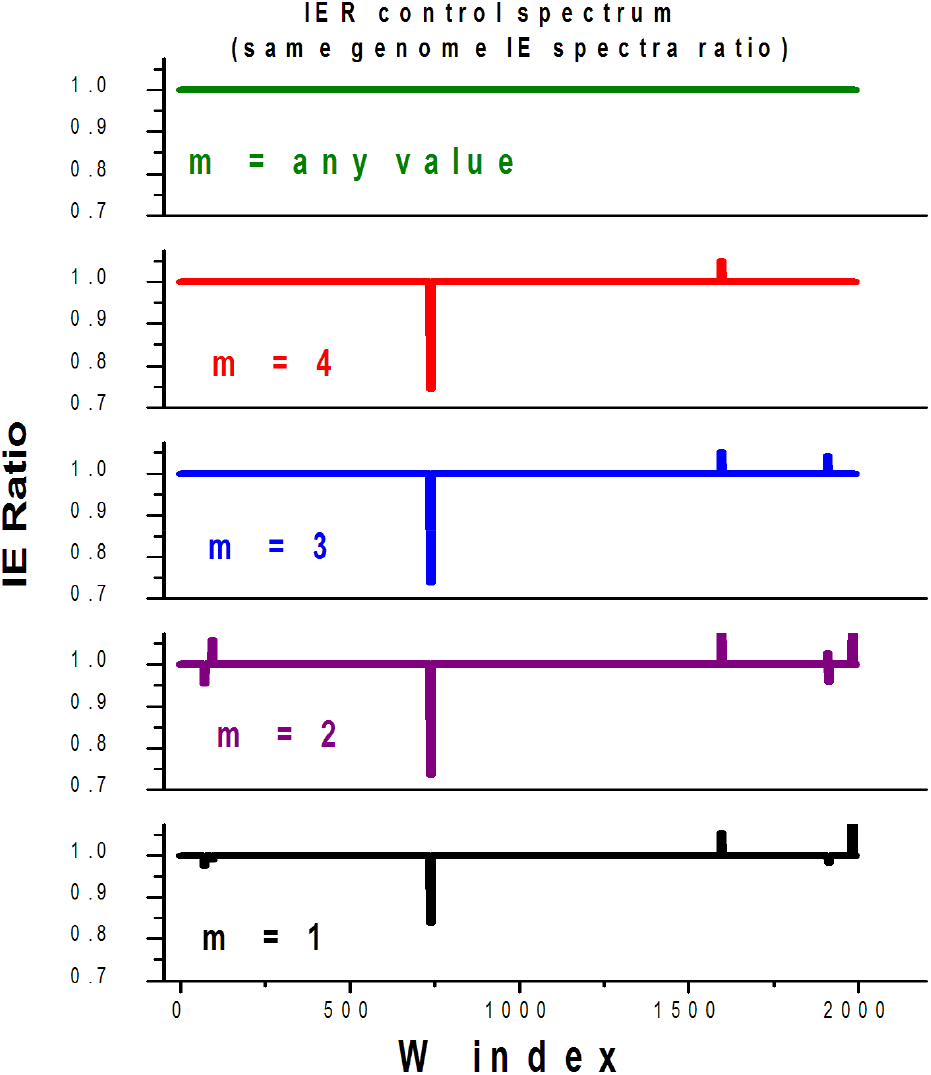
Information entropy ratio (IER) spectra of SARS-CoV-2 reference sequence (MN908947) to its variant (MT370518), produced using the GENIES software for WS = SS = 15, with variable m-block size. Top spectrum that shows no mutations is a control spectrum obtained by running the same reference genome on the software, while the rest show the IE ratio of the Wuhan reference sequence to its Taiwan variant.

## 5. Conclusions

We proposed to use the information theory to study genome sequences by creating the information entropy (IE) spectra of the genomes. When two or more genomes of the same family are analyzed in this way, one of them being considered as a reference sequence and the others as mutated versions, we showed that the ratio of the IE spectra, called the information entropy ratio (IER) spectra can be used to successfully identify genetic mutations in the analyzed sequences. This methodology requires computer automation and we produced GENIES, a stand-alone computer program, which is freely available to the scientific community [26]. GENIES is a fully functional code, that has an easy to use graphical interface and allows maximum versatility in choosing the computational parameters such as SS, WS and m-block size. However, the program needs further improvements, as presently it can only compare same size sequences, detecting single base point mutations, but excluding insertions and deletions.

The proposed methodology and the program’s functionality have been demonstrated on the SARS-CoV-2 reference sequence from Wuhan (MN908947) and one of its variants collected in Taiwan (MT370518). Both sequences have 29903 base points, but the program can analyze any kind of DNA / RNA genome sequence, of any size. Our results indicate that the best choice of the window size is 9 < WS ≤ 33, and the most optimal m-block size is m = 2, as this successfully captured all known mutations in our SARS-CoV-2 test sequences. While m = 2 is a generally applicable rule to the study of any sequence, the optimal WS values determined here for SARS-CoV-2 might be different for other sequences of different sizes. Besides block entropy approach, other complex indices [21, 35] have been proposed for detection or study of mutations, and these could be incorporated in the future versions of the GENIES tool. However, the true potential of this proposed methodology is achieved when using it in reverse, as a potential predictor of future genetic mutations. The idea is to analyze the inflection points where known mutations occurred, allowing to corroborate *special features* in the IE spectrum to the index location of the mutations, that in turn would work as a predictor of future genetic mutations. This is beyond the scope of this article, but we hope that this work will stimulate future studies based on GENIES and the proposed information entropy spectrum methodology.

## Acknowledgements

MV is grateful for the support to this work received from the School of Mathematics and Physics, University of Portsmouth. SR is part funded from Research England’s Expanding Excellence in England (E3) Fund.

## References

1. C.E. Shannon, A mathematical theory of communication, The Bell System Technical Journal, Vol. 27, pp. 379–423 (1948)

2. J. Machta, Entropy, information, and computation, American Journal of Physics 67:12, 1074–1077 (1999); https://doi.org/10.1119/1.19085

3. M. Hellman, “An extension of the Shannon theory approach to cryptography,” in IEEE Transactions on Information Theory, vol. 23, no. 3, pp. 289–294, May 1977, doi: 10.1109/TIT.1977.1055709.

4. P.M. Woodward, I.L. Davies, Information theory and inverse probability in telecommunication, Proceedings of the IEE - Part III: Radio and Communication Engineering (1952), 99 (58) : 37 http://dx.doi.org/10.1049/pi-3.1952.0011

5. L. Stark, G.C. Theodoridis, 2 - Information Theory in Physiology, Engineering Principles in Physiology, Pages 13-32, (1973) Academic Press, ISBN 9780121362010, https://doi.org/10.1016/B978-0-12-136201-0.50009-X

6. S. Naranan, V.K. Balasubrahmanyan, Information theoretic models in statistical linguistics - Part I: A model for word frequencies. Current Science, 63: 261–269 (1992).

7. F. Francesco, Shannon information theory and molecular biology, Journal of Interdisciplinary Mathematics, 12:1, 41–87, (2009) doi: 10.1080/09720502.2009.10700611

8. S. Yufeng, J. Fengxiang, Landslide Stability Analysis Based on Generalized Information Entropy, International Conference on Environmental Science and Information Application Technology, Wuhan, (2009), pp. 83-85, doi: 10.1109/ESIAT.2009.258.

9. A. Rhee, R. Cheong, A. Levchenko, The application of information theory to biochemical signaling systems, Phys. Biol. 9 045011 (2012) https://doi.org/10.1088/1478-3975/9/4/045011

10. Roman S. Ingarden, Quantum information theory, Reports on Mathematical Physics, Volume 10, Issue 1, 1976, Pages 43-72, ISSN 0034-4877, https://doi.org/10.1016/0034-4877(76)90005-7

11. Christian Beck, Generalised information and entropy measures in physics, Contemporary Physics, 50:4, 495–510, (2009) doi: 10.1080/00107510902823517

12. M.M. Vopson, The mass-energy-information equivalence principle, AIP Adv. 9, 095206 (2019).

13. T.A. Reichert, D.N. Cohen, A.K.C. Wong, An application of information theory to genetic mutations and the matching of polypeptide sequences, J. Theoret. Biol. 42, 245–261 (1973).

14. C. Cosmi, V. Cuomo, M. Ragosta, M.F. Macchiato, Characterization of nucleotide sequences using maximum entropy techniques, J. Theoret. Biol. 147, 423–432 (1990).

15. H. Herzel, W. Ebeling, A.O. Schmitt, Entropies of biosequences: The role of repeats, Phys. Rev. E 50, 5061–5071 (1994).

16. W. Li, K. Kaneko, Long-range correlations and partial 1/f spectrum in a noncoding DNA sequence, Europhys. Lett. 17(7), 655–660 (1992).

17. C.K. Peng, S.V. Buldyrev, A.L. Goldberger, S. Havlin, F. Sciortino, M. Simon, H.E. Stanley, Long-range correlations in nucleotide sequences, Nature 356, 168–170 (1992).

18. L. Wentian, G.M. Thomas, K. Kunihiko, Understanding long-range correlations in DNA sequences, Physica D: Nonlinear Phenomena, Volume 75, Issues 1–3, 392–416 (1994) https://doi.org/10.1016/0167-2789(94)90294-1.

19. R. Roman-Roldan, P. Bernaola-Galván, J. Oliver, Application of information theory to DNA sequence analysis: A review, Pattern Recognition, Volume 29, Issue 7, (1996) https://doi.org/10.1016/0031-3203(95)00145-X.

20. A. Hariri, B. Weber, J. Olmsted III, On the validity of Shannon-information calculations for molecular biological sequences, J. Theoret. Biol. 147, 235–254 (1988).

21. S. Vinga, Information theory applications for biological sequence analysis, Briefings in Bioinformatics, vol. 15 (3) 376–389 (2014).

22. J. A. Tenreiro Machado, Shannon Entropy Analysis of the Genome Code, Mathematical Problems in Engineering, Article ID 132625 (2012) https://doi.org/10.1155/2012/132625

23. F. Fernandes, A.T. Freitas, J.S. Almeida, S. Vinga, Entropic Profiler – detection of conservation in genomes using information theory, BMC Research Notes, 2:72 (2009) doi:10.1186/1756-0500-2-72

24. J.A. Tenreiro Machado, António C. Costa, Maria Dulce Quelhas, Shannon, Rényie and Tsallis entropy analysis of DNA using phase plane, Nonlinear Analysis: Real World Applications, Volume 12, Issue 6, 3135–3144 (2011) https://doi.org/10.1016/j.nonrwa.2011.05.013.

25. A. Thomas, S. Barriere, L. Broseus, J. Brooke, C. Lorenzi, J.P. Villemin, G. Beurier, R. Sabatier, C. Reynes, A. Mancheron, W. Ritchie, GECKO is a genetic algorithm to classify and explore high throughput sequencing data, Commun. Biol. 2, 222 (2019). https://doi.org/10.1038/s42003-019-0456-9

26. GENIES software free download: https://sourceforge.net/projects/information-entropy-spectrum/

27. A.O. Schmitt, H. Herzel, Estimating the Entropy of DNA Sequences, Journal of Theoretical Biology, Vol. 188 (3), 369–377 (1997) https://doi.org/10.1006/jtbi.1997.0493.

28. Liò P, Politi A, Buiatti M, Ruffo S. High statistics block entropy measures of DNA sequences, J Theor Biol.; 180(2): 151–60 (1996) https://doi.org/10.1006/jtbi.1996.0091

29. S.S. Melnik, O.V. Usatenko, Entropy and long-range correlations in DNA sequences, Computational biology and chemistry, Volume 53, Part A, 26–31 (2014) https://doi.org/10.1016/j.compbiolchem.2014.08.006

30. Genetic Information Entropy Spectrum (GENIES) User manual, 10 December (2020), DOI: 10.13140/RG.2.2.36557.46569

31. F. Wu, S. Zhao, B. Yu, Y.M. Chen, W. Wang, Z.G. Song, Y. Hu, Z.W. Tao, J.H. Tian, Y.Y. Pei, M.L. Yuan, Y.L. Zhang,F.H. Dai, Y. Liu, Q.M. Wang, J.J. Zheng, L. Xu, E.C. Holmes, Y.Z. Zhang, A new coronavirus associated with human respiratory disease in China, Nature 579, 265–269 (2020)

32. https://www.ncbi.nlm.nih.gov/nuccore/MN908947

33. D. Thanos, W. Li, A. Provata, Entropic fluctuations in DNA sequences, Physica A: Statistical Mechanics and its Applications, Vol. 493, 444–457, (2018) https://doi.org/10.1016/j.physa.2017.11.119.

34. https://www.ncbi.nlm.nih.gov/nuccore/MT370518

35. A. Provata, C. Nicolis, G. Nicolis, DNA viewed as an out-of-equilibrium structure, Phys. Rev. E 89, 052105 (2014) https://doi.org/10.1103/PhysRevE.89.052105

